# The impact of worldwide, national and sub-national severity distributions in Burden of Disease studies: a case study of individual cancer types in Scotland

**DOI:** 10.1101/654327

**Authors:** Grant MA Wyper, Ian Grant, Eilidh Fletcher, Gerry McCartney, Diane L Stockton

## Abstract

**Aim:** The main aim of this study was to consider the extent to which the use of worldwide severity distributions in Burden of Disease studies are influencing cross-country comparisons, by comparing Global Burden of Disease distributions with nationally derived severity distributions in Scotland for cancer types.

**Methods:** We obtained individual records from the Scottish Cancer Registry for 21 cancer types and linked these to registered deaths. We estimated prevalent cancer cases for 2016 and assigned each case to sequelae using Global Burden of Disease (GBD) 2016 study definitions. We compared the impact of using severity distributions based on GBD 2016, a Scotland-wide distribution, and a distribution specific to deprivation strata in Scotland, on the weighted-average disability weights for each cancer type in Scotland.

**Results:** The relative difference in point estimates of weighted-average disability weights based on GBD 2016 worldwide severity distributions compared with Scottish national severity distributions resulted in overestimates in the majority of cancers (17 out of 21 cancer types). The largest overestimates were for gallbladder and biliary tract cancer (70.8%), oesophageal cancer (31.6%) and pancreatic cancer (31.2%). Furthermore, the use of weighted-average disability weights based on Scottish national severity distributions rather than sub-national Scottish severity distributions stratified by deprivation quintile overestimated weighted-average disability weights in the least deprived areas (16 out of 18 cancer types), and underestimated in the most deprived areas (16 out of 18 cancer types).

**Conclusion:** Our findings illustrate a bias in point estimates of weighted-average disability weights created using worldwide severity distributions. This bias would have led to the misrepresentation of non-fatal estimates of the burden of individual cancers, and underestimated the scale of socioeconomic inequality in this non-fatal burden. This highlights the importance of not interpreting non-fatal estimates of burden of disease too precisely, especially for sub-national estimates and those comparing populations when relying on data inputs from other countries. It is essential to ensure that any estimates are based upon the best available country-specific data at the lowest granularity.

## Introduction

Burden of Disease (BOD) studies include both morbidity and mortality by framing them in terms of health loss suffered as a function of time [1]. Estimates of the frequency of morbidity in a population, such as prevalence, are transformed into Years Lived with Disability (YLD) using disability weights (DW) for each disease-specific sequelae. The proportional distribution of sequelae within a disease is commonly referred to as the severity distribution [2]. Recent advances in the Global Burden of Disease (GBD) study has seen the number of disease sequelae triple from 1,160 in GBD 2010 [3] to 3,484 in GBD 2017 [4]. While advances in the development of the granularity of disease and sequelae have been made, they have not been matched by the development of country-specific severity distributions [2]. This has prompted the development of country-specific severity distributions [5–7], although the impact of using these in favour of GBD severity distributions has not yet been evaluated.

There is a growing appetite for BOD studies to provide more granular estimates for regions within countries in both the GBD study [8–10] and independent national studies [5, 6, 11], to influence local policy development. This raises further questions about the utility of worldwide severity distributions in sub-national calculations. Findings from the GBD UK study raised some concerns over the accuracy of local estimates of YLD, noting that country-specific electronic health records have a role to play in refining estimates [8]. Whilst electronic health records can inform better disease estimates, it is important to advocate for and develop better prevalence estimates at the disease sequelae level to remove the reliance of worldwide severity distributions. Disability associated with different levels of sequelae varies significantly and thus have a major impact on YLD estimates. In short, if we assume a fixed proportion of prevalence in each sequelae within a disease across countries or in sub-populations within a country, we risk introducing a systematic bias in the YLD results.

The main aim of this study was to consider the extent to which worldwide severity distributions are influencing cross-country comparisons, by comparing them with nationally derived severity distributions in Scotland across a range of cancer types. A secondary aim was to investigate the impact of sub-national severity distributions compared to nationally derived severity distributions.

## Methods

### Data

There were 21 cancer types included in this study. Details of the cancer types included and excluded in this study are available in the Supplementary Appendix. Cancer types were restricted to those that had four common sequelae: (i) diagnosis and primary therapy phase; (ii) controlled phase; (iii) metastatic phase; and (iv) terminal phase. This approach was chosen to avoid attribution of differences due to interpretation of the GBD 2016 model when dealing with additional specific sequelae, such as procedural. DWs for each of the four sequelae of cancer were derived from the GBD 2016 study [12]. Unadjusted DWs were used to illustrate the theoretical scale of effect. The worldwide prevalence of the sequelae of each cancer type for were sourced from GBD 2016 [13] and apportioned into a severity distribution.

Scottish national and sub-national severity distributions were derived using individual patient records from the Scottish Cancer Registry, which holds registration records from 1980 onwards of all incident cancers diagnosed within the NHS in Scotland [14]. The disease model used to define the sequelae of each case was developed using definitions from the GBD 2016 technical appendix [15]. This involved calculating a 10-year prevalence of the incidence cohort for each cancer type to establish prevalent cases for 2016. Prevalent cases were apportioned to each sequelae using fixed durations found in the GBD 2016 technical appendix for the diagnosis and primary therapy phase, metastatic phase and terminal phase. Cases were assigned to the controlled phase if they did not satisfy the time-based criteria of the other three sequelae.

Patients were followed up over time and the date and cause of death were obtained from the National Records of Scotland’s register of deaths [16]. Using deterministic matching [17] of a patient-identifier between the cancer registry and register of death, we could confidently classify and exclude cases in accordance with the fixed duration cancer survival model definitions for each cancer type [15]. An area-based deprivation score was available for each patient, defined by the Scottish Index of Multiple Deprivation 2016 [18], which allowed us to create severity distributions for patients living in the most deprived fifth of areas, and those living in the least deprived fifth of areas in Scotland. The same fixed durations were used to classify sequelae across all methods.

### Analyses

Point estimates of the weighted-average DW for each cancer type were calculated based on four different severity distributions scenarios: (i) GBD 2016 worldwide; (ii) Scotland overall; (iii) the most deprived fifth of local areas in Scotland; and (iv) the least deprived fifth of local areas in Scotland. Relative and absolute differences in the point estimate of weighted-average DWs were assessed between approaches (i) and (ii) as the primary outcome. Absolute differences were presented by rescaling the DW into days of health loss by multiplying the DW by 365.25 days (where the additional 0.25 of a day takes into account the occurrence of a leap year every four years).

In addition, relative and absolute differences between (ii) and (iii), and (ii) and (iv) were assessed as secondary outcomes. The secondary analyses were restricted to 18 cancer types. Three cancer types (gallbladder and biliary tract, mesothelioma and nasopharynx) were excluded in the sub-national analyses because their deprivation-stratified severity distributions were based on a total number of prevalent cases less than 100.

### Data permissions and access

Formal permission to access linked National Health Service (NHS) administrative databases was granted by the Privacy Advisory Committee, NHS National Services Scotland (NSS) [PAC Reference 51/14] [19]. Patient-identifiable data extracts were extracted by NHS NSS and provided to NHS Health Scotland in the form of aggregate statistics that were subsequently used in this study. All summary data used in this study are provided in the supplementary appendix.

## Results

### GBD 2016 worldwide compared to Scottish national severity distributions

The relative difference across cancer types between the point estimate of the weighted-average DW based on GBD 2016 worldwide severity distributions compared with Scottish national severity distributions resulted in positive values across 17 out of 21 cancer types (Figure 1). In these 17 cancer types, using GBD 2016 severity distributions would have resulted in a relative overestimate of the point estimate of the weighted-average DW. The largest relative overestimates were observed for: gallbladder and biliary tract cancer (70.8%), oesophageal cancer (31.6%) and pancreatic cancer (31.2%). There were four instances where the use of GBD 2016 severity distributions would have resulted in a relative underestimate of the point estimate of the weighted-average DW: mesothelioma (−20.9%), brain and nervous system cancer (−7.1%), uterine cancer (−4.8%) and thyroid cancer (− 2.4%).

**Figure 1:**
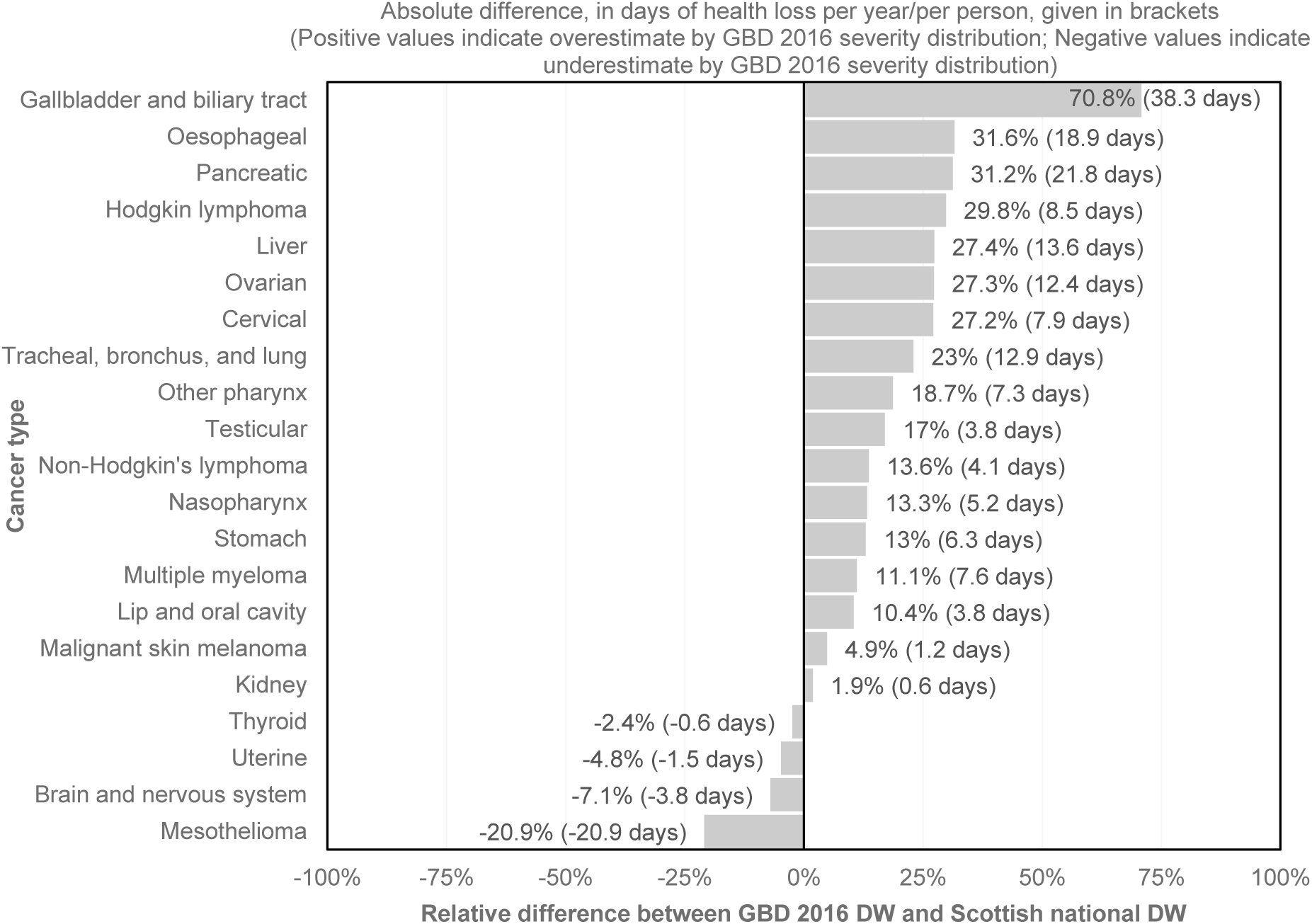
Relative and absolute comparison of cancer disability weights: GBD 2016 worldwide versus Scottish national

In terms of absolute difference, the largest overestimates would have been made for gallbladder and biliary tract cancer (38.3 additional days of health loss per year, per person), pancreatic cancer (21.7 additional days of health loss per year, per person) and oesophageal cancer (18.9 additional days of health loss per year, per person). The largest absolute underestimate would have been made for mesothelioma (20.9 fewer days of health loss per year, per person).

### Scottish national compared to sub-national severity distributions

When the relative difference between the point estimates of the weighted-average DW based on Scottish national severity distributions was assessed against severity distributions based on the most deprived fifth of Scottish areas, there were negative values in 16 out of 18 cancer types (Figure 2). In these 16 cancer types, using nationally derived severity distributions would have resulted in a relative underestimate of the point estimate of the weighted-average DW. The largest relative underestimates were observed for: lip and oral cavity cancer (−11.6%), oesophageal cancer (−8.4%) and other pharynx cancer (−8.1%). There were two instances where the use of Scottish national severity distributions would have resulted in a relative overestimate of the point estimate of the weighted-average DW: brain and nervous system cancer (10.3%), and Hodgkin lymphoma (0.1%).

**Figure 2:**
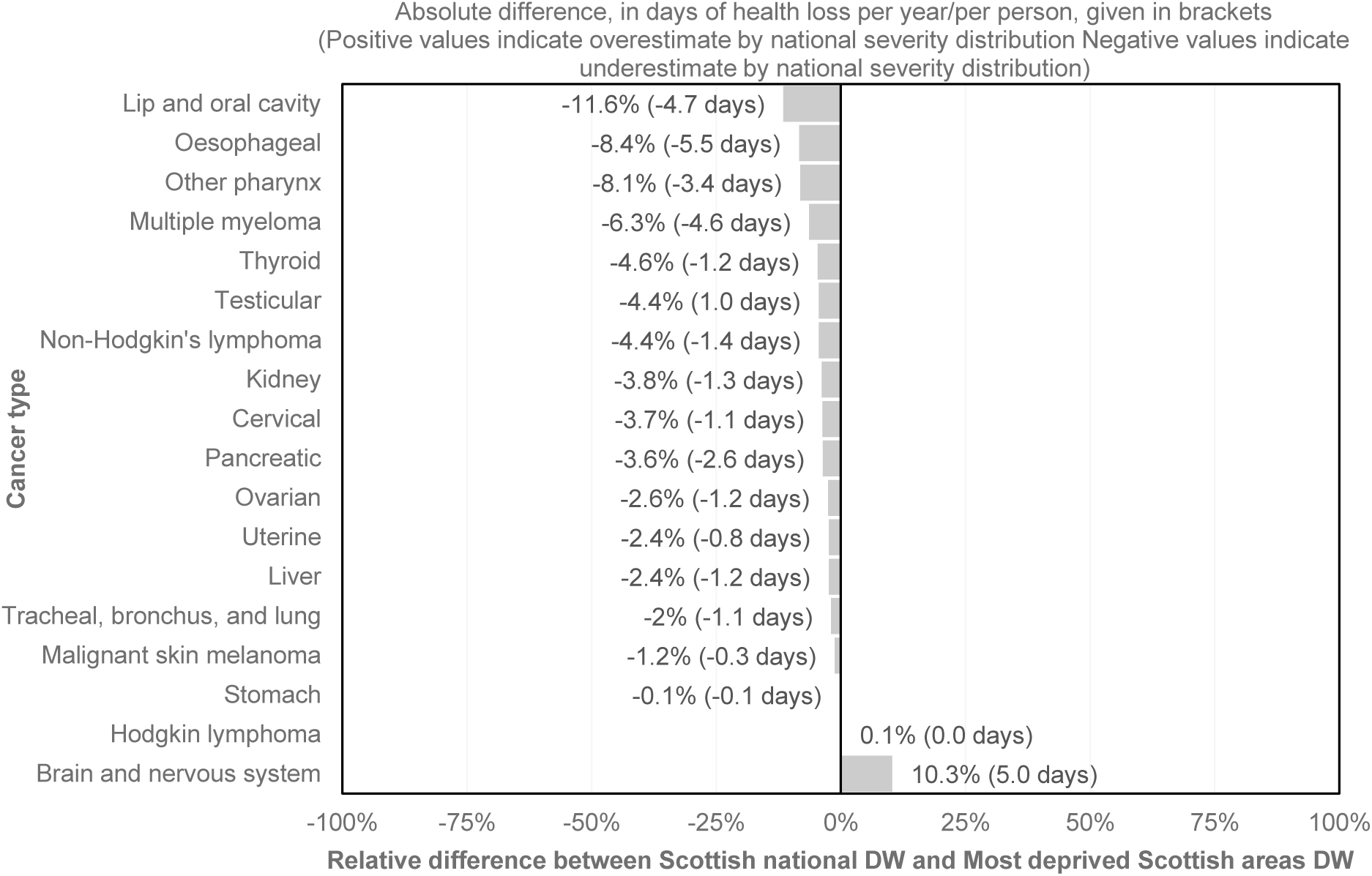
Relative and absolute comparison of cancer disability weights: Scottish national versus most deprived fifth of Scottish areas

The cancer types where the largest absolute underestimates would have been made were for oesophageal cancer (5.5 fewer days of health loss per year, per person), lip and oral cavity cancer (4.7 fewer days of health loss per year, per person) and multiple myeloma (4.6 fewer days of health loss per year, per person). The largest absolute overestimate would have been made for brain and nervous system cancer (5.0 additional days of health loss per year, per person).

The relative difference across cancer types between the point estimate of the weighted-average DW based on Scottish national severity distributions compared to severity distributions based on the least deprived fifth of Scottish areas resulted in positive values across 16 out of 18 cancer types (Figure 3). In these 16 cancer types, using nationally derived severity distributions would have resulted in an overestimate of the weighted-average DW. The largest relative overestimates were observed for: other pharynx cancer (16.9%), pancreatic cancer (9.9%) and lip and oral cavity cancer (8.7%).

**Figure 3:**
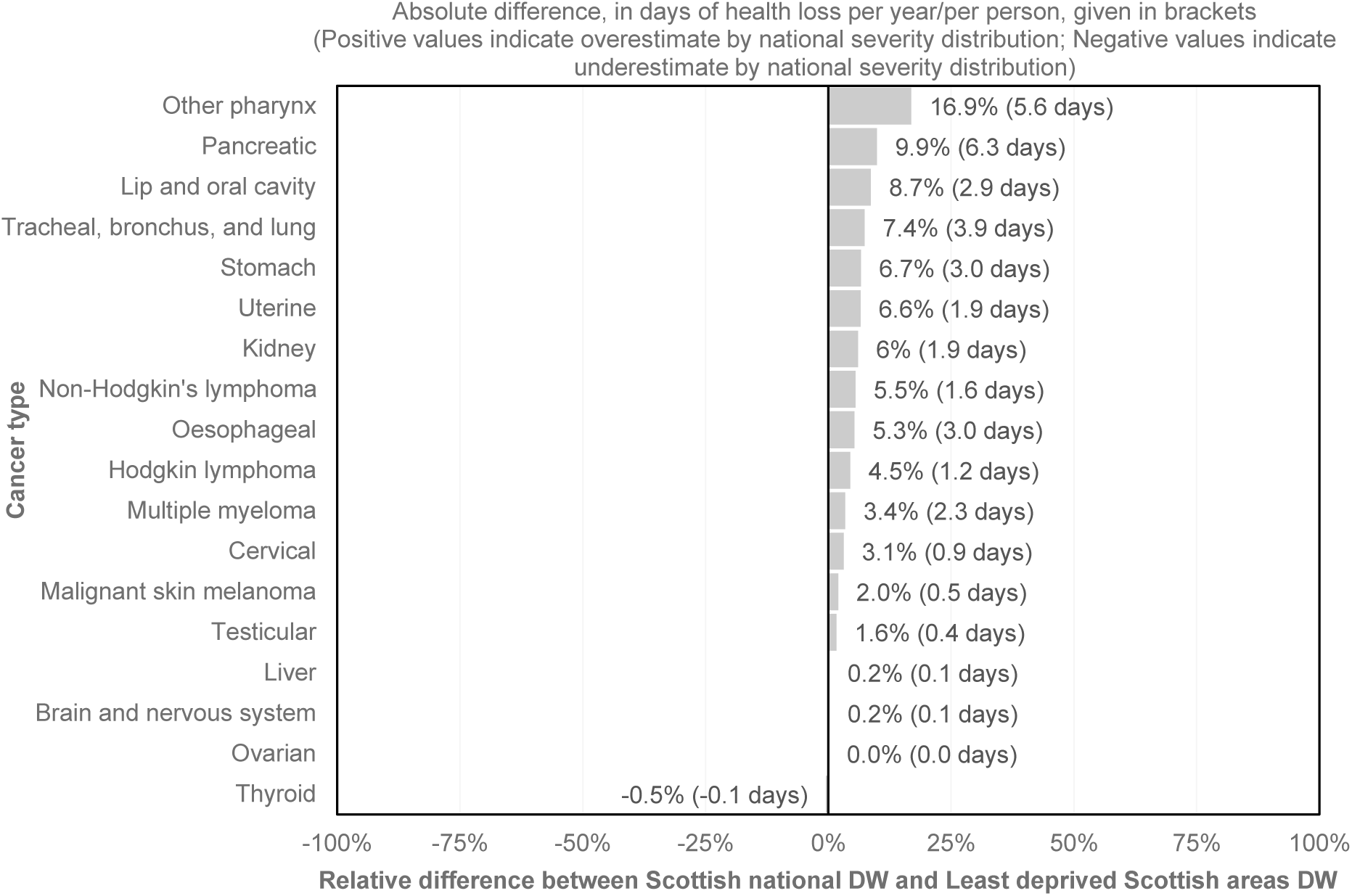
Relative and absolute comparison of cancer disability weights: Scottish national versus least deprived fifth of Scottish areas

The largest absolute differences in overestimation were observed for pancreatic cancer (6.3 additional days of health loss per year, per person), other pharynx cancer (5.6 additional days of health loss per year, per person) and tracheal, bronchus and lung cancer (3.9 additional days of health loss per year, per person).

The magnitude of the differences in point estimates of weighted-average DWs were much larger when comparing GBD 2016 worldwide with nationally derived severity distributions, than when comparing differences between national and sub-national severity distributions.

## Discussion

### Summary of findings

This study is the first to evaluate the impact of using published GBD 2016 worldwide, compared to national and sub-national derived severity distributions on weighted-average disability weights. Our study has illustrated that the use of GBD 2016 worldwide severity distributions would have led to overestimation in the point estimate of disability weights assigned to individual cancer types in a large majority of cases. The range of overestimation varied in size, with the largest relative overestimates being observed in gallbladder and biliary tract cancer (70.8%), oesophageal cancer (31.6%) and pancreatic cancer (31.2%). Although it was less common, there were some instances whereby using GBD 2016 worldwide severity distributions would have resulted in underestimation. This was observed for mesothelioma (−20.9%), brain and nervous system cancer (−7.1%) and uterine cancer (− 4.8%).

Additionally, we have illustrated the importance in considering differences in severity distributions across sub-regions of a country. Our assessment of sub-national severity distributions stratified by area-based deprivation quintiles indicated that the use of Scottish national severity distributions would have led to an underestimate of the point estimate of the weighted-average disability weight in the most deprived areas of Scotland across the majority of cancer types. Conversely, the use of Scottish national severity distributions would have led to overestimates of point estimates of weighted-average disability weights in the least deprived areas of Scotland. Considering both these findings, using national severity distributions would have understated the extent of socioeconomic inequalities in YLD associated with individual cancer types.

### How this compares with existing literature

Existing studies of BOD severity distributions are limited. Those already published have been designed to develop severity distributions, and have not evaluated the impact of using different severity distributions [2, 7]. A study in Korea estimated severity distributions for eight diseases using two independent national surveys [7]. Both methods resulted in similar patterns of severity across all eight diseases. The GBD severity distributions study acknowledges concerns over applying estimates of severity distributions based on data from the United States and Australia, noting that it is the only available information that they were able to use [2].

### Strengths and weaknesses

A major strength of this study lies within the use of the Scottish Cancer Registry [13] with patient-linked death registrations records [15, 16]. This allowed us to precisely classify each incident case to an exact sequelae. The availability of a patient postcode of residence on each cancer registration record allowed us to classify each case to an area-based deprivation quintile. The transparency of the GBD 2016 models appendix [14] allowed us to utilise the definitions and allocate cases on a like-for-like basis. Whilst the use of patient-linked electronic health records is a major strength in our study, we acknowledge that for many countries it would be impractical to obtain estimates of severity for sub-strata of the population, or even nationally due to lack of data availability, access permissions or small population size [20].

We have chosen to produce sub-national severity distributions based on deprivation, as our secondary focus was around local area estimates with a focus on socioeconomic inequalities. We also acknowledge the limitation that deprivation is not the solely significant factor and that on a disease by disease basis, other stratifications such as ethnicity or gender may be more appropriate and display important intersectional effects [21]. In addition, we also note that our study has only assessed one of the aspects of potential error and bias in estimates; severity distributions. There are other aspects, such as the disability weight assigned to a particular health state that could also bias estimates. At present the same four disability weights are used across each cancer sequelae. Further research is needed to assess whether these estimates are appropriate, or whether disability weights across cancers, and indeed other causes, are context-specific and socially influenced [22]. We have applied the same cancer survival across our comparisons. Given there is a reasonable likelihood that survival times would indeed vary across and within countries, our findings on the scale of difference are a likely to be a conservative account.

### Implications for research and policy

This study has important implications for policy and practice. In highlighting potential biases in the over and underestimation of point estimates of disability weights, we have illustrated a proportional bias that will carry through to estimates of YLD. In terms of cancer, the magnitude of bias on Disability-Adjusted Life Years (DALYs) would be minor because the majority of DALYs are generated through Years of Life Lost to premature mortality (YLL). However the bias remains for cancer YLD estimates, and these concerns extend to other causes of disease, particularly leading causes of YLD and causes of YLD that exhibit wide inequalities [23–26].

Further research is required to understand the impact of severity distributions for conditions that may have larger inequalities in case fatality, or severity, than cancers to explore differences across a larger range of diseases. Caution should be taken over interpreting the applicability of our severity distributions to other countries due to differences in demography, social circumstance and access to healthcare systems across countries. There are large resource requirements in undertaking BOD studies. Both independent BOD researchers and collaborators of the GBD need to know where to focus their efforts in order to obtain the maximum benefit of information. Our results have illustrated the importance of ensuring that any estimates are based upon the best available country-specific data at the lowest possible level, in the context of cancer types.

BOD studies are a means to influence policy and practice, and these findings are important in highlighting a systematic bias in the point estimates that are being used to rank causes that is largely being overlooked. To a degree, the lack of sub-national severity distributions would negate each other in national results, but would mask differences between regions that have heterogeneous levels of deprivation and therefore underestimate the true extent of inequalities. Care must be taken in interpreting YLD estimates too precisely, especially with sub-national estimates and estimates comparing populations.

## Acknowledgements

We would like to acknowledge the GBD 2016 cancer collaborators for making detailed and transparent model definitions publically available as part of the GBD 2016 technical appendix. Additionally, we would like to thank members of the European Commission funded ‘Joint Action on Health Information (InfAct)’ workshop in Paris in April 2019 for productive discussions on the use of severity distributions in BOD studies.

